## Supplementary information file for "Use of Raman and Raman optical activity to extract atomistic details of saccharides in aqueous solution"

#### 1 Methods

##### 1.1 Target Saccharides

Four disaccharides (α-D-glucopyranosyl-(1→1)-α-D-glucopyranose (trehalose), methyl 1α-2α-mannobiose (M12), methyl 1α-3α-mannobiose (M13), and methyl 1α-6α-mannobiose (M16)) were studied to investigate the effects of rotation around glycosidic bonds ($\phi_1/\phi_2/\phi_3$) on Raman/ROA spectra. The glycosidic angles are depicted in Figure 1 and explicitly defined in Table 1.

Table 1: Atom definitions of dihedral angles associated with glycosidic links of studied trehalose, M12, M13, and M16 disaccharides shown in Figure 1.

| | $\phi_1$ | $\phi_2$ | $\phi_3$ |
|---|---|---|---|
| trehalose | H1-C1-O1'-C1' | C1-O1'-C1'-H1' | - |
| M12 | H1-C1-O2'-C2' | C1-O2'-C2'-H2' | - |
| M13 | H1-C1-O3'-C3' | C1-O3'-C3'-H3' | - |
| M16 | H1-C1-O6'-C6' | C1-O6'-C6'-C5' | O6'-C6'-C5'-H5' |

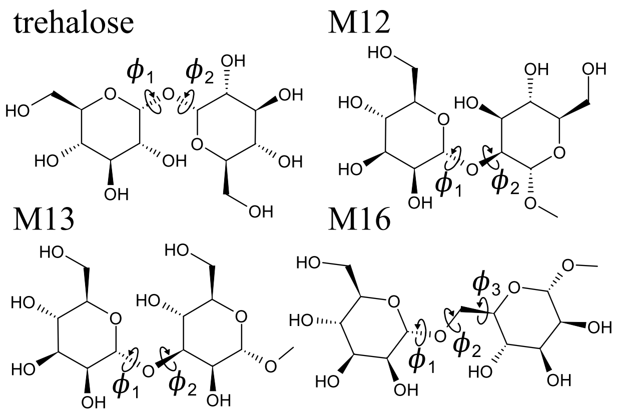


Figure 1: Investigated disaccharides: trehalose, methyl-1α-2α-mannobiose (M12), methyl-1α-3α-mannobiose (M13), and methyl-1α-6α-mannobiose (M16), together with defined glycosidic dihedral angles $\phi_1$, $\phi_2$, and $\phi_3$ (atom definitions in Table 1).

#### 1.2 Computational methods

Simulation of Raman/ROA spectra and connection to experimental data consist of several steps. 1) A representative ensemble of structures is generated using molecular dynamics simulations. 2) Raman/ROA spectra are calculated for each of the structures using quantum chemical methods, obtaining ensemble-averaged spectra. 3) We score the similarity of simulated spectra with experiment, and if applicable, we find the best fit of simulated data to the Raman/ROA spectra. 4) If possible, we compare our calculations to NMR data. Underneath, we explain each of these steps.

##### 1.2.1 Ensemble Generation

Molecular dynamics (MD) simulations were used to sample the phase space and to generate representative ensembles used to simulate spectra of studied saccharides. At first, we describe the general setup of MD simulations, then we go through MD simulations, and finally, we describe additional details to individual MD simulations.

**General MD Setup and Simulation Summary.** All MD simulations were performed using the Gromacs-2016-4[1] simulation package patched with Plumed 2.5.[2] All studied system were prepared in a sufficiently large simulation box to accomodate a single saccharide solvated with water molecules(e.g., 3×3×3 nm or 3.5×3.5×3.5 nm box sizes resulting into a single saccharide and 800-1500 water molecules). The only exception was the crowding effects study, where two monosaccharides were placed in a simulation box instead of one. Since all of the studied saccharides are expected to be mainly present in a $^4C_1$ chair conformation,[3] we use it as a default puckering conformation. Both α and β anomers of Glc, GlcA, GlcNAc were considered. In our MD simulations we used glycam-6h[4] (sugars) and OPC3[5] water force fields. Simulations were performed in an NpT ensemble using 2 fs time step, where pressure was handled using the Parrinello-Rahman[6] barostat at 1 bar with the couping constant of 5 $ps^{-1}$. Temperature was controlled using the Nose-Hoover thermostat[7] at 300 K using a time constant 1 $ps^{-1}$. Long range electrostatic interactions were modeled using PME[8] with a 1 nm cutoff for the real part. Van der Waals interactions were also treated with 1 nm cutoff and the long range dispersion correction for energy and pressure was applied.[9] Both electrostatic and Van der Waals interactions were shifted to zero at the cutoff. The neighbour searching was done using Verlet list. All hydrogen bonds containing hydrogen were constrained using LINCS.[10] Rest of specific MD simulation details for each of the application is covered in the next sections.

Generally, we performed three types of calculations. The first (unbiased MD, see Table 2) is an unbiased MD simulation from which representative ensemble of structures was obtained, for which subsequently Raman/ROA spectra was calculated. The second type of calculation ('free energy') is well-tempered metadynamics MD simulation[11] to find a free energy profile along studied coordinate such as glycosidic dihedral angles or puckering coordinates. We aimed to find local minima on given free energy profile for which we performed the third type of a simulation. The third type of a simulation (biased MD) consists of restrained system in found local minima using harmonic position restrains. Ensemble average spectra for these regions were then calculated. The minima are denoted *md*1, *md*2, *md*3, *md*4, *md*5, and *md*6. The ensemble averaged spectra were then used to best fit experimental data to estimate contributing weights to the overall spectra. The representative ensembles were obtained by uniformly sampling the performed MD simulations. As a default we extracted 250 structures, with exception of when studying Binary Mixtures and Crowding Effects, where 500 structure have been extracted. Table 2 summarizes all performed simulations in this work.

Table 2: Summary of MD simulations performed in this work.

| **Performed Simulations** | |
|---|---|
| **Conformation of Glycosidic Bonds (trehalose, M12, M13, and M16)** | |
| unbiased MD | 500 ns |
| free energy profile | metadynamics MD in $\phi_1/\phi_2/\phi_3$ glycosidic dihedral angles |
| biased MD | *md*1/*md*2/*md*3/*md*4/*md*5/*md*6 conformers 200 ns |
| **Probing Puckering Conformations (MeGlcA)** | |
| unbiased MD | 500 ns/250 structures |
| free energy profile | metadynamics MD in puckering coordinates |
| biased MD | $^1C_4$/ $^4C_1$/$^OS_2$/$^1S_3$ puckering conformers 50 ns |
| **α/β anomeric equilibrium constant (Glc, GlcA, and GlcNAc)** | |
| unbiased MD | 500 ns for each of the α/β anomer |
| **Raffinose Trisaccharide** | |
| unbiased MD | 500 ns |
| **Binary Mixtures** | |
| unbiased MD | 500 ns for pure MeGlc and MeGlcNAc |
| **Crowding Effects** | |
| unbiased MD | 500 ns for a single MeGlc |
| unbiased MD | 500 ns for two MeGlc in close proximity[a] |

Further details on individual MD simulations follow.

**Conformation of Glycosidic Bonds.** A 200 ns well tempered metadynamics MD in $\phi_1/\phi_2/\phi_3$ dihedral angles was performed to obtained corresponding free energy profiles. Gaussians with a height of 6 kJ and width 0.35 rad were deployed every 250 steps with a biasfactor 12($\Delta$T = 3300K). We found that the free energy profile of M16 in $\phi_3$ shows little variability, see Section S2.1) and therefore we integrated it out, obtaining 2D free energy profiles in $\phi_1/\phi_2$ glycosidic dihedral angles for all four investigated disaccharides. We sampled the local minima (up to six local minima) on the free energy profile using a harmonic position restrains with a force constant k = 50 kJ.mol$^{-1}$.nm$^{-2}$. Since all of the studied disaccharides are expected to occur predominantly in the $^4$C$_1$ chair conformation,[3] we restricted pyranose rings of both sugar monomers in all studied disaccharides to the $^4C_1$ puckering conformation. This task was performed using the Plumed upper walls restraint, where we set both $\theta$ puckering coordinates of both rings to $<$0.75.[2]

**Probing Puckering Conformation.** A well-tempered metadynamics MD of methyl-β-D-glucuronic acid was used to calculate the free energy profile in $q_x$, $q_y$, and $q_z$ puckering coordinates on a grid of 100×100×100 points spaced equidistantly in a range of $(-0.1, 0.1)nm$. The MD simulation was performed for 50 ns, where gaussians of height 1 kJ and width 0.01 nm were deployed every 250 steps with a bias factor 6($\Delta$T = 1500K). Afterward, all phase space points on the grid were converted to populations through the Boltzmann factor, and cartesian coordinates were transformed to radial ($Q$, $\phi$, and $\theta$). Since all occupied phase space points essentially lie on the surface of a sphere, it is practical to integrate the populations (points) in the $Q$ space with a radial cutoff of $0.05 \cdot \pi$ around given ($\phi$,$\theta$) point. Integrated populations were then converted back to free energies and placed on the free energy surface, which we show in Mollweide equal-area projection, where the free energies are finally interpolated to obtain a smooth free energy surface. The advantage of this approach is a free energy surface, where two phase space points can be directly compared since they were obtained from equal phase space volumes.

From the puckering free energy profile, we sampled four local minima (labeled as $^1C_4$, $^4C_1$, $^OS_2$, and $^1S_3$) in a 50 ns biased MD simulation using the $\phi$ and $\theta$ puckering coordinates. The systems were restrained using harmonic restraint with a force constant of k = 50 kJ.mol$^{-1}$.rad$^{-2}$. Since $^1C_4$ and $^4C_1$ puckering conformers are located on the poles of the puckering free energy profile, we restrained these only in $\theta$ puckering variable. Table 3 summarizes restrain values that were used to sample $^1C_4$/ $^4C_1$/$^OS_2$/$^1S_3$ conformers.

Table 3: Restrain values [rad] of $\phi/\theta$ puckering coordinates used in biased MD simulations of methyl-β-D-glucuronic acid yielding $^1C_4$/ $^4C_1$/$^OS_2$/$^1S_3$ conformers. $^1C_4$ and $^4C_1$ conformer were restrained only in $\theta$ puckering variable.

| | $^1C_4$ | $^4C_1$ | $^OS_2$ | $^1S_3$ |
|---|---|---|---|---|
| MeGlcA | all/3.14 | all/0 | 3.14/1.57 | 1.8/1.57 |

##### 1.2.2 Calculation of Raman/ROA Spectra

After extracting individual snapshots from the MD trajectory, we proceed to partial QM optimization, followed by calculation of vibrational frequencies and the Raman and ROA intensities. For this, we use a previously developed `hybrid` simulation approach.[12] In brief, this simulation protocol is a QM/MM based method that consists of extracting snapshots from the MD trajectory, where the central molecule together with surrounding water molecules (3 Å cutoff) are kept. The water molecules are treated at the MM level of theory, while the solute is calculated at the B3LYP/6-311++G** level of theory using the ONIOM method with electrostatic embedding.[13,14] TIP3P[15] and glycam-6h[4] force field parameters were used to describe the MM part of the ONIOM method. The long-range solvation effects are taken care of by using the COSMO implicit solvation model. After preparation, the system is optimized in ten steps unrestrained optimization. After optimization, we use the far-from-resonance coupled-perturbed Kohn-Sham theory[16,17] at 532 nm (SCP180) and harmonic approximation to calculate vibrational frequencies and Raman/ROA intensities. All calculated vibrational frequencies are rescaled using a frequency scaling function

$\phi(\tilde{\nu}, a, b, c, d) = \phi(\tilde{\nu}, 0.982, 1.00, 15, 1210)$.[12] Calculated intensities were corrected for the temperature factor at 300 K and convoluted using Lorentzian with a full width at half maximum (FWHM) $\Gamma = 7.5$ cm$^{-1}$. Full equation for calculation of averaged Raman/ROA spectra with already scaled vibrational frequencies $I^{sim}_{Raman/ROA}(\tilde{\nu})$ stands as

$$I^{sim}_{Raman/ROA}(\tilde{\nu}) = \frac{1}{\tilde{\nu}} \cdot \frac{1}{1 - e^{-\frac{\hbar \cdot \tilde{\nu}}{k_B T}}} \sum_{j=1}^{\#snapshots} \frac{\sum_{i=1}^{\#vib} I_{Raman/ROA,ij}(\tilde{\nu}_{ij}) \cdot \frac{2}{\pi} \cdot \frac{\Gamma}{4(\tilde{\nu}-\tilde{\nu}_{ij})^2+\Gamma^2}}{\#\ sim}, \quad (1)$$

For more details, refer to former work in the literature.[12] All the QM/MM calculations are performed using the Gaussian16 program package.[18]

**Spectra Quality Assessment.** We used the overlap integral $S$ defined in Eq 2 for qualitative comparison of simulated spectra with experiment.

$$S = \frac{\int_{\tilde{\nu}_{min}}^{\tilde{\nu}_{max}} I_{sim}(\tilde{\nu}) I_{exp}(\tilde{\nu}) d\tilde{\nu}}{\sqrt{\int_{\tilde{\nu}_{min}}^{\tilde{\nu}_{max}} {I_{sim}}^2(\tilde{\nu}) d\tilde{\nu} \int_{\tilde{\nu}_{min}}^{\tilde{\nu}_{max}} {I_{exp}}^2(\tilde{\nu}) d\tilde{\nu}}}, \quad (2)$$

where $I_{sim}$ and $I_{exp}$ represent the simulated and experimental spectra that are to be compared. In the case of the Raman spectrum, the integral $S$ provides values between 0 and 1, where values above 0.95 are considered as a very good agreement. For the ROA spectra, $S$ can yield values between -1 and 1, with -1 being obtained for exact mirror images. Values above 0.7 are considered as a good agreement for the ROA spectrum.

**The Best Fit to Experimental Raman/ROA Data.** The experimental spectra is a cumulative average of all spectra produced by any accessible state by the system during the experiment. Computationally, we search for states ($i$) that contribute significantly to the overall simulated spectra ($I^{sim}_{Raman}$ and $I^{sim}_{ROA}$). Importantly for this work, each simulated state ($I^{sim}_{i,Raman}$ and $I^{sim}_{i,ROA}$) can represent either different molecular stereochemistries (e.g., spectra of α/β anomer forms), or conformational states (e.g., different puckering conformers) which are molecular features that will be discussed in this work. The overall spectra are weighted averages of each considered spectra, $I^{sim}_{i,Raman}$ and $I^{sim}_{i,ROA}$:

$$I^{sim}_{Raman}(\vec{A}) = \sum_{i=0}^{spectra} A_i \cdot I^{sim}_{i,Raman},$$
$$I^{sim}_{ROA}(\vec{A}) = \sum_{i=0}^{spectra} A_i \cdot I^{sim}_{i,ROA}. \tag{3}$$

where $A_i$ is the weight given to a specific state $i$ and $\vec{A}$ is the vector of weights for each state ($\{A_i\}$). These vector represents the relative presence of each of the states in the real experimental ensemble. Therefore, the same $A_i$ weights applies the Raman and ROA spectra of a given state $i$. A simulated state contributes significantly to the experimental spectra when its calculated spectra is required to better fit the experimental spectra. Finding the best agreement between weighted simulation spectra and experimental data is performed by minimizing the following cost function:

$$F(\vec{A}) = (1 - S_{Raman}(\vec{A}))^2 + (1 - S_{ROA}(\vec{A}))^2, \tag{4}$$

where $S_{Raman}(\vec{A})$ and $S_{ROA}(\vec{A})$ are overlap integrals defined in Eq.2.

In this work, instead of optimizing/weighting directly from single states, we calculate first an intermediate average spectra of a phase space region of interest. These are what we call local simulated spectra ($I^{sim}_{i,Raman}$ and $I^{sim}_{i,ROA}$) and they are a simple ensemble average spectra of every simulated state corresponding to the region of interest.

**Estimation of Uncertainty of Determined Weights.** Fitting experimental data with preselected structures results in one unique solution (the global minimum). However, the question is how robust this solution is, i.e., whether there are other solutions with very similar spectra but different conformer distributions. To investigate how robust is our fit, we numerically calculate ranges for the abundance of each of the conformers that result at least into 99 % of the found global minimum in terms of the sum of the overlap integrals (i.e., $S_{Raman} + S_{ROA}$). This reveals the conformer weights' ranges that results to essentially the same final simulation spectra as the best fit solution (i.e.,

$S_{Raman} + S_{ROA} > 0.99(S_{Raman}^{bestfit} + S_{ROA}^{bestfit})$). The spread of the weights values $\{A_i^{min}, A_i^{max}\}$ represents the uncertainty of obtained conformer weights and determines the accuracy of the method to determine the real weight of a particular conformer/feature.

**Obtaining NMR $^3J$ Spin-spin Coupling Constants.** We have also estimated abundance of states sampled by biased MD using NMR $^3J$ spin-spin coupling constants. We calculated spin-spin coupling constants either by means of QM chemistry (i.e., glycosidic bonds of M12 and M13 disaccharides and puckering of methyl-β-glucuronic acid), or using the Karplus equation[19] (i.e., M12 and M13 disaccharides). The ensemble average $^3J$ spin-spin couplings of sampled conformers were used to best fit the experimental data as described below.

To calculate QM derived coupling constants we started with structures that were already optimized as described in the section 1.2.2. Optimized structures were stripped of all water molecules, leaving only the sugar moiety. Then, the spin-spin coupling constants were calculated with the Gaussian program package[18] using the mPW1PW91/pc-J2[20] level of theory, together with the CPCM continuum solvation. Only the Fermi contact terms were considered. Alternatively, vicinal $^3J$ were calculated using the Karplus equation with corresponding dihedral angles as described in Ref.[19]

Similarly as for Raman/ROA, we can fit experimental NMR spin-spin coupling constants ($^3J_i^{exp}$). We follow the same procedure as with Raman/ROA data using a cost function $G(\vec{A})$ defined as

$$G(\vec{A}) = \sum_{i=0}^{coupling\ constants} ({}^3J_i^{sim} - {}^3J_i^{exp})^2, \tag{5}$$

where $^3J_i^{sim}$ is the weighted simulation spin-spin coupling constant and is calculated as

$$^3J_i^{sim} = \sum_{j=0}^{conformers} A_j \cdot {}^3J_j^{sim}. \tag{6}$$

$^3J_j^{sim}$ represents the spin-spin coupling constant of a conformer obtained from averaging individual snapshots (i.e., states) within a region. $A_j$ represents the weight of a given conformer. Minimizing the cost function yields the best fit to experimental data, i.e., optimized weights $\vec{A}$. We estimated the uncertainty of determined conformer weights similarly as in the Raman/ROA calculations. We numerically found the ranges for the abundance of each of the conformers that would lead to a maximum deviation of 0.2 Hz from the best fit solution, which is the reported experimental error.[19,21] Within the error range all spin-spin couplings have to satisfy condition $|^3J_i^{sim} -^3 J_i^{sim,bestfit}| < 0.2$.

#### 2 Secondary structure of trehalose and methyl-1α-2α-mannobiose (M12) disaccharides

Figure S2 shows calculated free energy profiles in $\phi_1/\phi_2$ dihedral angles (left). Middle panel shows 250 structures that were extracted from unbiased 500 ns MD simulation on top of the free energy profiles. Right panel then shows structures which were extracted from biased MD simulations using harmonic position restraints at given phase space point yielding $md1/md2/md3/md4$ conformers(restrain values in Table S4).

Table 4: Restrain values [rad] of $\phi_1/\phi_2$ glycosidic angles used in biased MD simulations of trehalose and M12 yielding $md1/md2/md3/md4$ conformers.

| | $md1$ | $md2$ | $md3$ | $md4$ |
|---|---|---|---|---|
| trehalose | -0.9/-0.9 | 1/-0.65 | 0.45/2.83 | 1.1/1.1 |
| M12 | -0.8/0.4 | 0.8/0.4 | -0.6/2.9 | -1.0/-1.0 |

Note that trehalose's free energy profile is symmetric (x axis) as the molecule is also symmetric, therefore we are probing all local minima.

Figures S3 and S4 then shows Raman/ROA ensemble averaged spectra of local-conformers (top), the best fit to the experimental data (top left), and spectra obtained using structures from MD simulation (top right).

In Table S5 we report the MD composition, together with the best fit and the error estimates for trehalose and methyl-1α-2α-mannobiose

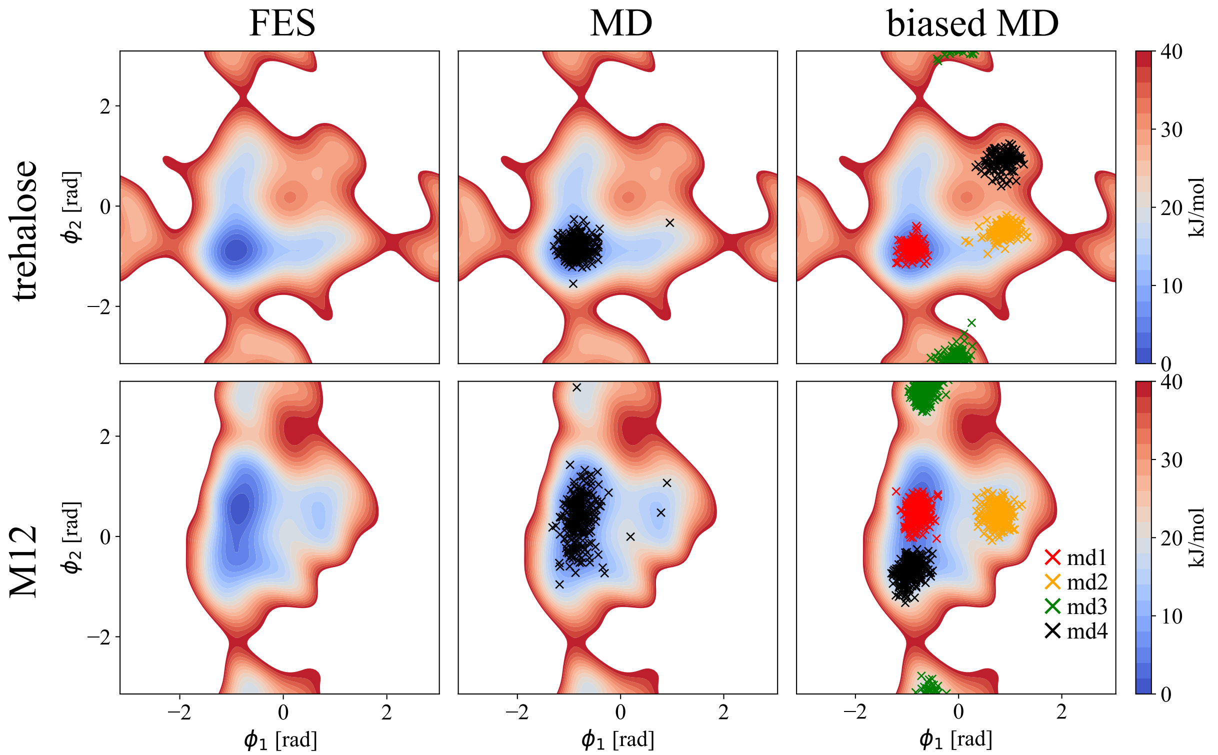


Figure 2: Left: Calculated free energy surface(FES) of studied disaccahrides in $\phi_1/\phi_2$ dihedral angles. Middle: Calculated FES, together with 250 extracted structures from unbiased 500 ns MD simulations (MD). Right: Calculated FES, together with 250 extracted structures per each biased 200 ns MD simulation(biased MD) (red - $md1$, orange - $md2$, green - $md3$, black - $md4$. White regions represents area where the free energy is larger than 40 kJ/mol.

Table 5: Abundance of conformers of trehalose and methyl-1α-2α-mannobiose (M12) as obtained by MD, NMR, and best fit to Raman/ROA (error bars in brackets).

| trehalose | MD | NMR | Best fit |
|---|---|---|---|
| md1 | 1.00 | 1.00[22] | 1.00 (0.92-1.00) |
| md2 | 0.00 | 0.00[22] | 0.00 (0.00-0.07) |
| md3 | 0.00 | 0.00[22] | 0.00 (0.00-0.08) |
| md4 | 0.00 | 0.00[22] | 0.00 (0.00-0.05) |
| M12 | | | |
| md1 | 0.84 | 0.95 (0.00-0.99)[b] | 0.86 (0.56-0.99) |
| md2 | 0.01 | 0.00 (0.00-0.13)[b] | 0.00 (0.00-0.14) |
| md3 | 0.00 | 0.00 (0.00-0.42)[b] | 0.00 (0.00-0.29) |
| md4 | 0.15 | 0.05 (0.00-0.65)[b] | 0.14 (0.00-0.40) |

*means of error estimation in SI
[b]exp data and Karplus equation[19]

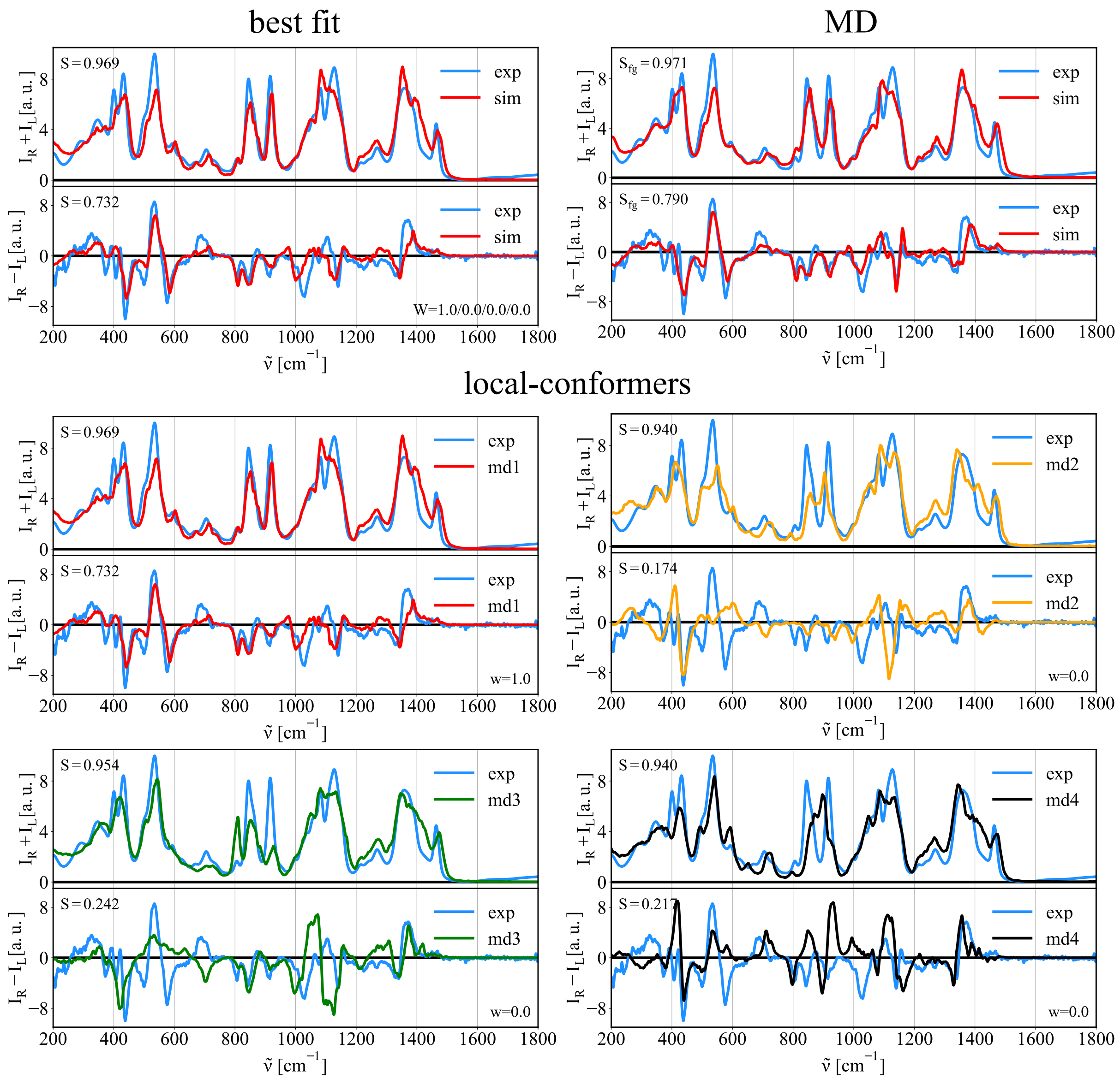


Figure 3: Trehalose: Top right - MD: Simulated Raman/ROA spectra of disaccharide obtained while using structures obtained from unbiased MD simulation (MD) and comparison to experimental data (exp). Bottom - local-conformers: Calculated ensemble averaged Raman and ROA spectra of disaccharide prepared in 4 distinct conformations $md1/md2/md3/md4$ as described in Figure 2 and comparison to experimental data. Top left - best fit: Best fit of $md1/md2/md3/md4$ Raman/ROA spectra to experimental data.

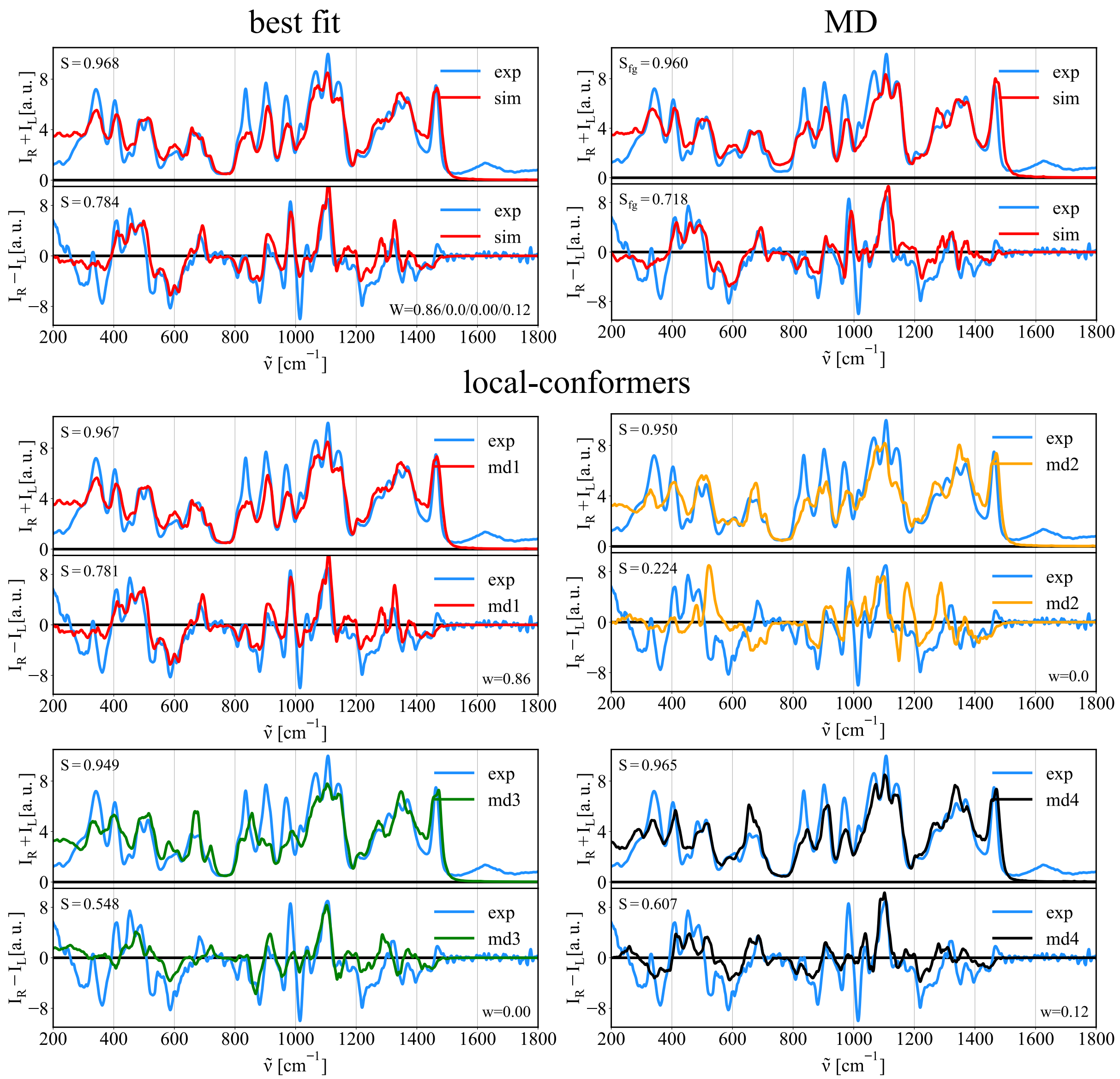


Figure 4: Methyl-1α-2α-mannobiose: Top right - MD: Simulated Raman/ROA spectra of disaccharide obtained while using structures obtained from unbiased MD simulation (MD) and comparison to experimental data (exp). Bottom - local-conformers: Calculated ensemble averaged Raman and ROA spectra of disaccahride prepared in 4 distinct conformations $md1/md2/md3/md4$ as described in Figure 2 and comparison to experimental data. Top left - best fit: Best fit of $md1/md2/md3/md4$ Raman/ROA spectra to experimental data.

##### 2.1 M16 $\phi_1/\phi_2/\phi_3$ FES

In Figure S5 we show that $\phi_3$ angle shows little variability and therefore, we integrated the variable out to obtain 2D free energy profile, i.e., FES in dihedral angles $\phi_1$ and $\phi_2$.

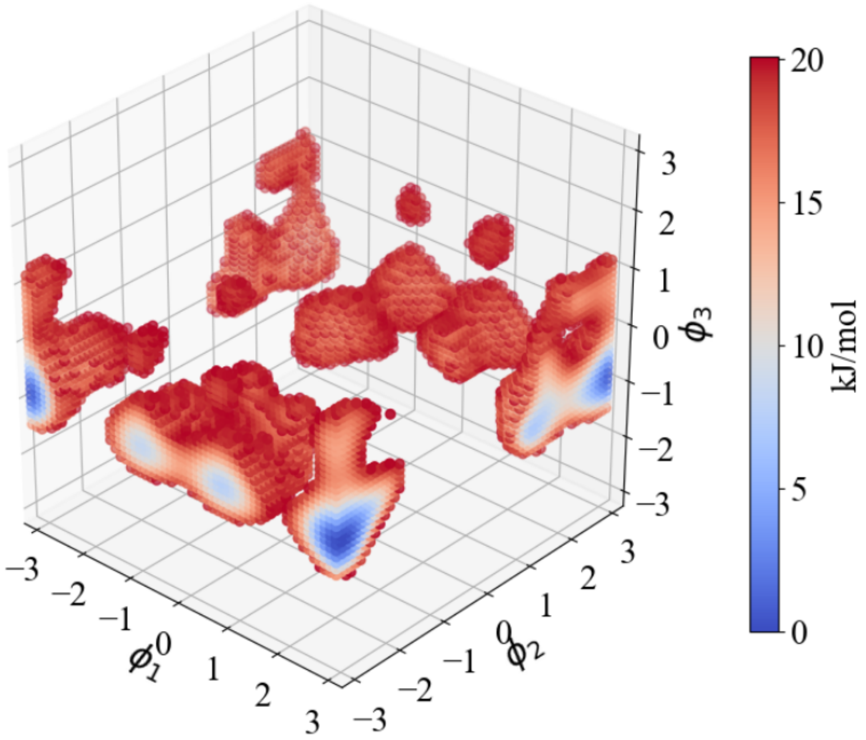


Figure 5: Calculated free energy profile of methyl-1α-6α-mannobiose in $\phi_1$, $\phi_2$, and $\phi_3$ dihedral angles showing a little variability in $\phi_3$. Therefore, $\phi_3$ variable was integrated out obtaining a 2D FES as shown in the main text.

##### 2.2 Calculation of NMR populations of local-conformers of M13 using QM calculations

Table S7 shows average calculated coupling constants for local-conformers of M13, together with the best fit and available experimental data.[23] Best fit weights are $md1/md2/md3/md4$ = 0.50/0.11/0.25/0.14.

Table 6: Calculated glycosidic spin-spin constants of M13 local-conformers using QM calculations together with the best fit and experimental data.

| | $md1$ | $md2$ | $md3$ | $md4$ | Best fit | Exp. data |
|---|---|---|---|---|---|---|
| M13($^3J_{H_1',C_3}$) | 4.57 | 2.92 | 4.04 | 2.48 | 3.80 | 3.80[23] |
| M13($^3J_{C_1',H_3}$) | 4.47 | 5.77 | 6.63 | 4.25 | 5.00 | 5.00[23] |

##### 2.3 Calculation of NMR populations of local-conformers of M12 and M13 using Karplus equation

Estimation of conformer populations of M12 and M13 from NMR data was pursued by means of using Karplus equation for $^3J_{CH}$ spin-spin coupling constants and MD data to obtain ensemble averaged simulation coupling constants $^3J_{CH}(sim)$ of local-conformers. The average calculated values were then used to obtain best fit to experimental data. Table S7 shows average calculated coupling constants for individual conformers, together with the best fit and available experimental data. Best fit weights are M12: $md1/md2/md3/md4$ = 0.95/0.00/0.00/0.05 and M13: $md1/md2/md3/md4$ = 0.62/0.20/0.16/0.01.

Table 7: Calculated glycosidic spin-spin constants of M12 and M13 local-conformers employing Karplus equation together with the best fit values and experimental data.

| - | $md1$ | $md2$ | $md3$ | $md4$ | Best fit | Exp. data |
|---|---|---|---|---|---|---|
| M12($^3J_{H_1',C_2}$) | 3.76 | 1.66 | 4.54 | 2.99 | 3.70 | 4.10[19] |
| M12($^3J_{C_1',H_2}$) | 4.91 | 5.40 | 6.96 | 3.71 | 4.85 | 4.60[19] |
| M13($^3J_{H_1',C_3}$) | 3.82 | 3.10 | 4.72 | 1.67 | 3.80 | 3.80[23] |
| M13($^3J_{C_1',H_3}$) | 4.61 | 4.96 | 6.56 | 5.05 | 5.00 | 5.00[23] |

### 3 Calculation of NMR populations of local-puckering-conformers of MeGlcA

Table S8 shows ensemble averaged calculated coupling constants for local-conformers, together with the best fit and available experimental data. Best fit weights for MeGlcA are

$^1C_4/^4C_1/^OS_2/^1S_3 = 0.01/0.99/0.00/0.00$.

Table 8: Calculated spin-spin constants of MeGlcA local-conformers employing using QM calculations together with the best fit values and experimental data.

| MeGlcA | $^1C_4$ | $^4C_1$ | $^OS_2$ | $^1S_3$ | Best fit | Exp. data[21] |
|---|---|---|---|---|---|---|
| $^3J_{12}$ | 1.20 | 7.83 | 0.71 | 0.60 | 7.77 | 8.80 |
| $^3J_{23}$ | 2.67 | 9.20 | 2.83 | 5.85 | 9.13 | 9.50 |
| $^3J_{34}$ | 2.83 | 8.95 | 0.87 | 10.93 | 8.88 | 8.50 |
| $^3J_{45}$ | 1.23 | 9.88 | 9.21 | 8.41 | 9.79 | 9.60 |

### 4 Raffinose trisaccharide

In this section, we report MD structural analysis of a raffinose trisaccharide. Reported populations were obtained from unbiased MD simulation (details main text). Additional, free energy surfaces (FES) were obtained using well-tempered metadynamics MD simulations, where only probed variables were biased. The simulation setups to obtain puckering FES of each pyranose were the same as described in the main text for monosaccharides. Similarly, the simulation setups for rotation around glycosidic angles were the same as for the disaccharides. Additionally, the puckering of fructose was also studied for which we used puckering coordinates for 5 membered rings (two phase space variables: the radial $Q$, and angular $\phi$). Similarly to 6 membered rings, $Q$ remains almost constant and only $\phi$ changes significantly in value. Therefore, we biased only $\phi$. To calculate FES of puckering of fructose, we deployed gaussians with height 0.1 kJ and $\sigma = 0.1$ every 250 steps while using biasfactor = 6. The simulation was run for 200 ns.

From Figure S6, we see that for the whole MD simulation the puckering of each sugar remains only in one state. Then, Figure S8 shows that during unbiased MD simulation the glycosidic angles between glucose and fructose remain almost invariant, while dihedral angles between galactose and glucose access several conformers. This behavior is similar to what was seen in case of methyl-1α-6α-mannobiose, a feature attributed to $1 \rightarrow 6$ glycosidic bonds.[24] The largest variations are observed in $\phi_1/\phi_2$ angles. The angle $\phi_3$ remains almost constant. In Figure S9 we calculated free energy surfaces for rotations around glycosidic angles between the monosaccharides. The results mimic the results from previous Figure S8, showing that in unbiased MD simulation do not miss any significant local minima and all relevant states are sampled.

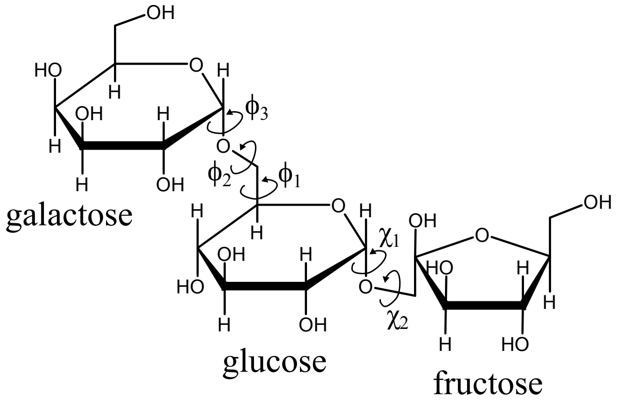


Figure 6: Raffinose trisacccharide: structure and investigated angles.

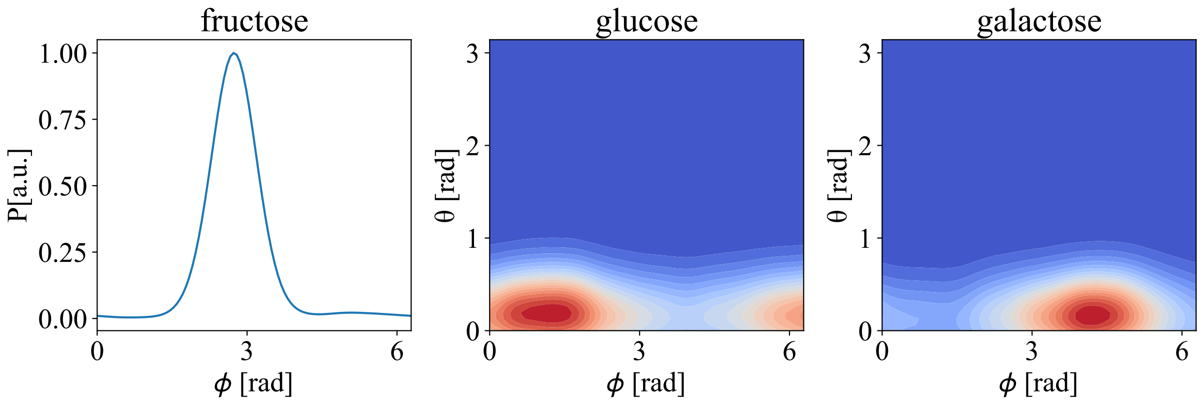


Figure 7: Raffinose trisacccharide: MD populations - puckering for each monosaccharide.

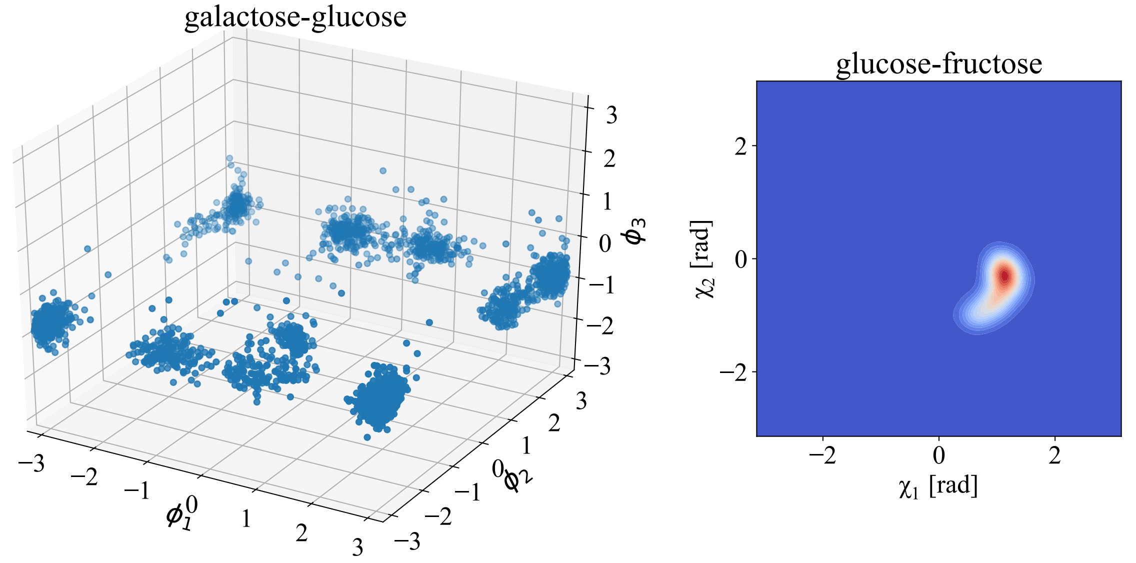


Figure 8: Raffinose trisacccharide: MD populations - glycosidic angles as defined in Figure 6

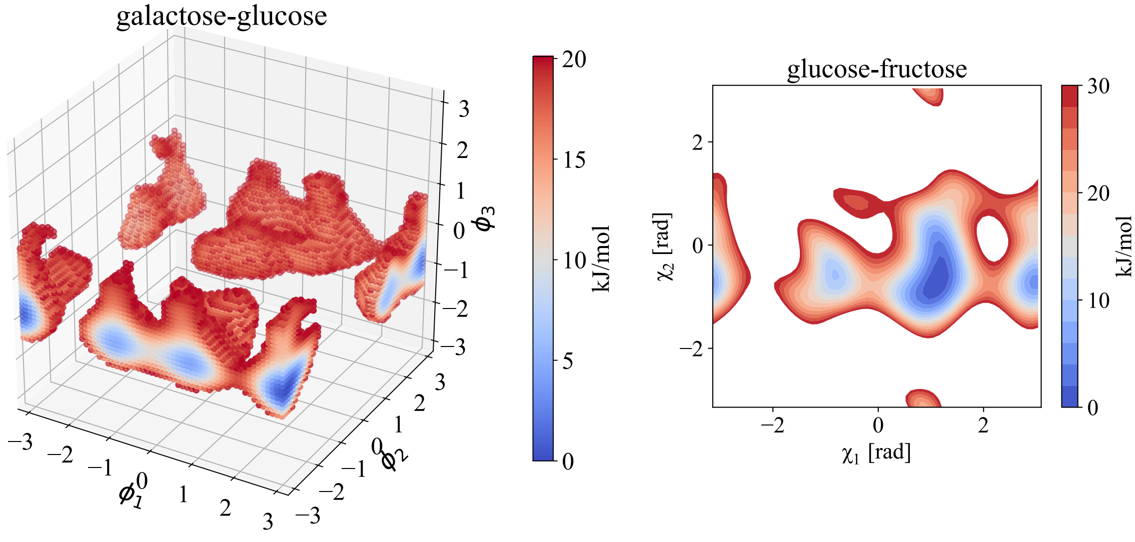


Figure 9: Raffinose trisacccharide: FES (obtained from metdynamics MD simulation) in glycosidic angles as defined in Figure 6. Empty regions represent regions with free energy > threshold (20/30 kJ/mol)

### 5 Binary mixture

Figure S10 shows calculated Raman/ROA spectra of pure substances (methyl-β-glucose (MeGlc) and methyl-β-N-glucosamine (MeGlcNAc)).

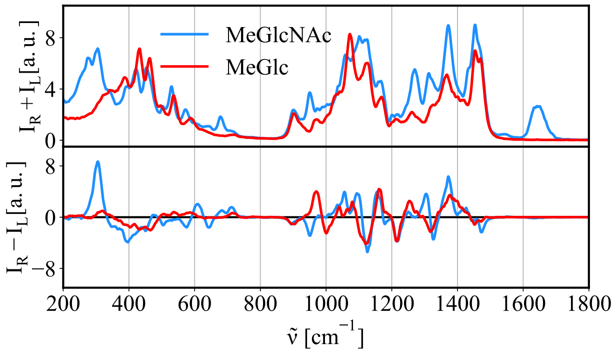


Figure 10: Simulated Raman/ROA spectra of methyl-β-glucose (MeGlc) and methyl-β-N-glucosamine (MeGlcNAc).

### 6 Crowding effects

Figure 11 shows that including empirical dispersion when studying effect of stacking has no effect on the outcome. Still, stacking has no effect on resulting Raman/ROA spectra.

(2) Bonomi, M.; Bussi, G.; Camilloni, C.; Tribello, G. A.; Banáš, P.; Barducci, A.; Bernetti, M.; Bolhuis, P. G.; Bottaro, S.; Branduardi, D.; Capelli, R.; Carloni, P.; Ceriotti, M.; Cesari, A.; Chen, H.; Chen, W.; Colizzi, F.; De, S.; De La Pierre, M.; Donadio, D.; Drobot, V.; Ensing, B.; Ferguson, A. L.; Filizola, M.; Fraser, J. S.; Fu, H.;

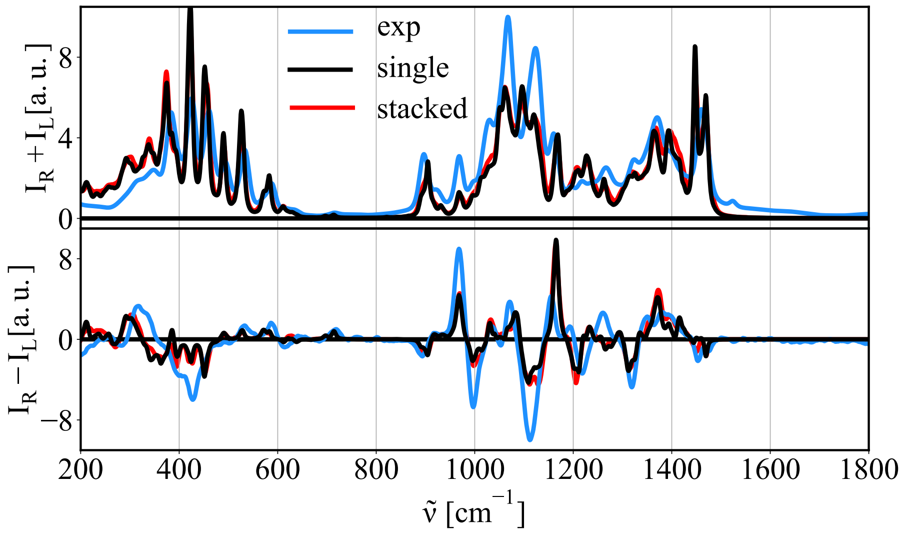


Figure 11: Calculated Raman and ROA spectra of methyl-β-glucose as in infinite dilution (black, a single molecule); while interacting with other methyl-β-glucose sugar (red), and experimental data (blue) for comparison as obtained using B3LYP/6-311++G** level of theory together with the CPCM solvation model with empirical dispersion corrections included (Grimme's D3 dispersion with Becke-Johnson damping). Prior to spectra calculation the systems were optimized in a 10 steps QM optimization.

Gasparotto, P.; Gervasio, F. L.; Giberti, F.; Gil-Ley, A.; Giorgino, T.; Heller, G. T.; Hocky, G. M.; Iannuzzi, M.; Invernizzi, M.; Jelfs, K. E.; Jussupow, A.; Kirilin, E.; Laio, A.; Limongelli, V.; Lindorff-Larsen, K.; Löhr, T.; Marinelli, F.; Martin-Samos, L.; Masetti, M.; Meyer, R.; Michaelides, A.; Molteni, C.; Morishita, T.; Nava, M.; Paissoni, C.; Papaleo, E.; Parrinello, M.; Pfaendtner, J.; Piaggi, P.; Piccini, G. M.; Pietropaolo, A.; Pietrucci, F.; Pipolo, S.; Provasi, D.; Quigley, D.; Raiteri, P.; Raniolo, S.; Rydzewski, J.; Salvalaglio, M.; Sosso, G. C.; Spiwok, V.; Šponer, J.; Swenson, D. W.; Tiwary, P.; Valsson, O.; Vendruscolo, M.; Voth, G. A.; White, A. Promoting transparency and reproducibility in enhanced molecular simulations. *Nature Methods* **2019**, *16*, 670–673.

(3) Sundararajan, P. R.; Rao, V. S. Theoretical studies on the conformation of aldopyranoses. *Tetrahedron* **1968**, *24*, 289–295.

(4) Kirschner, K. N.; Yongye, A. B.; Tschampel, S. M.; González-Outeiriño, J.;

Daniels, C. R.; Foley, B. L.; Woods, R. J. GLYCAM06: A generalizable biomolecular force field. Carbohydrates. *Journal of Computational Chemistry* **2008**, *29*, 622–655.

(16) Nafie, L. *Vibrational optical activity: Principles and applications*; Wiley: Chichester, 2011.

(17) Ruud, K.; Helgaker, T.; Bouř, P. Gauge-origin independent density-functional theory calculations of vibrational Raman optical activity. *Journal of Physical Chemistry A* **2002**, *106*, 7448–7455.

(18) Frisch, M. J.; Trucks, G. W.; Schlegel, H. B.; Scuseria, G. E.; Robb, M. A.; Cheeseman, J. R.; Scalmani, G.; Barone, V.; Petersson, G. A.; Nakatsuji, H.; Li, X.; Caricato, M.; Marenich, A. V.; Bloino, J.; Janesko, B. G.; Gomperts, R.; Mennucci, B.; Hratchian, H. P.; Ortiz, J. V.; Izmaylov, A. F.; Sonnenberg, J. L.; Williams-Young, D.; Ding, F.; Lipparini, F.; Egidi, F.; Goings, J.; Peng, B.; Petrone, A.; Henderson, T.; Ranasinghe, D.; Zakrzewski, V. G.; Gao, J.; Rega, N.; Zheng, G.; Liang, W.; Hada, M.; Ehara, M.; Toyota, K.; Fukuda, R.; Hasegawa, J.; Ishida, M.; Nakajima, T.; Honda, Y.; Kitao, O.; Nakai, H.; Vreven, T.; Throssell, K.; Montgomery Jr., J. A.; Peralta, J. E.; Ogliaro, F.; Bearpark, M. J.; Heyd, J. J.; Brothers, E. N.; Kudin, K. N.; Staroverov, V. N.; Keith, T. A.; Kobayashi, R.; Normand, J.; Raghavachari, K.; Rendell, A. P.; Burant, J. C.; Iyengar, S. S.; Tomasi, J.; Cossi, M.; Millam, J. M.; Klene, M.; Adamo, C.; Cammi, R.; Ochterski, J. W.; Martin, R. L.; Morokuma, K.; Farkas, O.; Foresman, J. B.; Fox, D. J. Gaussian16 Revision B.01. 2016.
